# Insights into platypus population structure and history from whole-genome sequencing

**DOI:** 10.1101/221481

**Authors:** Hilary C. Martin, Elizabeth M. Batty, Julie Hussin, Portia Westall, Tasman Daish, Stephen Kolomyjec, Paolo Piazza, Rory Bowden, Margaret Hawkins, Tom Grant, Craig Moritz, Frank Grutzner, Jaime Gongora, Peter Donnelly

## Abstract

The platypus is an egg-laying mammal which, alongside the echidna, occupies a unique place in the mammalian phylogenetic tree. Despite widespread interest in its unusual biology, little is known about its population structure or recent evolutionary history. To provide new insights into the dispersal and demographic history of this iconic species, we sequenced the genomes of 57 platypuses from across the whole species range in eastern mainland Australia and Tasmania. Using a highly-improved reference genome, we called over 6.7M SNPs, providing an informative genetic data set for population analyses. Our results show very strong population structure in the platypus, with our sampling locations corresponding to discrete groupings between which there is no evidence for recent gene flow. Genome-wide data allowed us to establish that 28 of the 57 sampled individuals had at least a third-degree relative amongst other samples from the same river, often taken at different times. Taking advantage of a sampled family quartet, we estimated the *de novo* mutation rate in the platypus at 7.0×10^−9^/bp/generation (95% CI 4.1×10^−9^ − 1.2×10^−8^/bp/generation). We estimated effective population sizes of ancestral populations and haplotype sharing between current groupings, and found evidence for bottlenecks and long-term population decline in multiple regions, and early divergence between populations in different regions. This study demonstrates the power of whole-genome sequencing for studying natural populations of an evolutionarily important species.

## Introduction

Next-generation sequencing technologies have greatly facilitated studies into the diversity and population structure of non-model organisms. For example, whole-genome sequencing (WGS) has been applied to investigate demographic history and levels of inbreeding in primates, with implications for conservation (Locke et al., 2011; Prado-Martinez et al., 2013; McManus et al., 2015; Xue et al., 2015; Abascal et al., 2016). It has also been used to study domesticated species such as pigs (Li et al., 2013; Bosse et al., 2014), dogs (Freedman et al., 2014), maize (Hufford et al., 2012) and bees (Wallberg et al., 2014), to infer the origins of domestication, its effect on effective population size (*N_e_*) and nucleotide diversity, and to identify genes under selection during this process. Some studies have identified signatures of introgression (Bosse et al., 2014) or admixture (Miller et al., 2012; Lamichhaney et al., 2015) between species, which is important to inform inference of past *N_e_*. Others have used WGS data to identify particular genomic regions contributing to evolutionarily important traits, such as beak shape in Darwin’s finches (Lamichhaney et al., 2015), mate choice in cichlid fish (Malinsky et al., 2015), and migratory behaviour in butterflies (Zhan et al., 2014). Here, we describe a population resequencing study of the platypus (*Ornithorhynchus anatinus*), which is one of the largest such studies of non-human mammals, and the first for a non-placental mammal.

In addition to laying eggs, platypuses have a unique set of characteristics (Grant, 2007), including webbed feet, a venomous spur (only in males), and a large bill that contains electroreceptors used for sensing their prey. Their karyotype is 2n = 52 (Bick and Sharman, 1975), and they have five different male-specific chromosomes (named Y chromosomes), and five different chromosomes present in one copy in males and two copies in females (X chromosomes), which form a multivalent chain in male meiosis (Grutzner et al., 2004).

Though apparently secure across much of its eastern Australian range, the platypus has the highest conservation priority ranking among mammals when considering phylogenetic distinctiveness (Isaac et al., 2007). Given concerns about the impact of climate change (Klamt et al., 2011), disease (Gust et al., 2009) and other factors on platypus populations, there is a need to better understand past responses of platypus populations to climate change, and the extent of connectivity across the species range.

The first platypus genome assembly (ornAna1) was generated using established whole-genome shotgun methods (Warren et al., 2008) from a female from the Barnard River in New South Wales (NSW) (see Figure 1). This assembly was highly fragmented and did not contain any sequence from the Y chromosomes.

**Figure 1:**
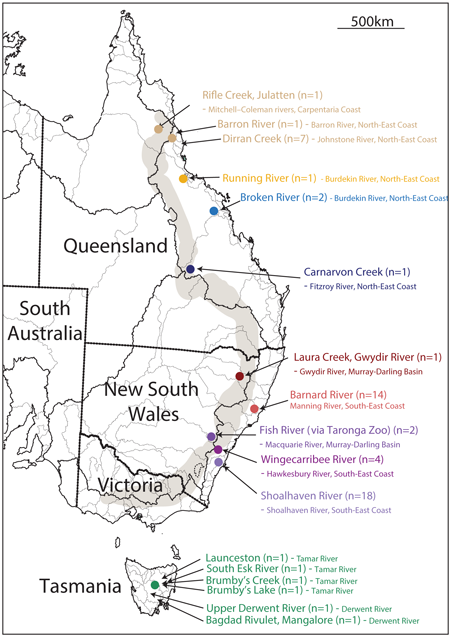
Map showing our sampling locations within river regions in eastern and southeastern Australia. The lighter lines on the figure are rivers and the darker lines demarcate catchment areas. The waterways our samples come from are indicated by arrows, with sample sizes in brackets after the river name. The specific catchments these rivers fall into and their corresponding larger drainage division are indicated in small letters after the river name. The text colours correspond to those used in later figures. The transparent grey region represents the Great Dividing Range (GDR). Note that all samples except those from the Fish River, Gwydir River and Rifle Creek come from river basins that drain east from the GDR. This map is adapted from one obtained from the Australian Bureau of Meteorology (www.bom.gov.au/water/geofabric/documents/BOM002_Map_Poster_A3_Web.pdf).

The initial genome paper included only a limited analysis of inter-individual variation and population structure based on 57 polymorphic retrotransposon loci. Subsequently, several other studies have investigated diversity and population structure using microsatellites or mitochondrial DNA (mtDNA) (Kolomyjec et al., 2009; Gongora et al., 2012; Furlan et al., 2013; Kolomyjec et al., 2013) (Table 1). They reported much stronger differences between than within river systems, but found some evidence of migration between rivers that were close together, implying limited overland dispersal.

**Table 1:** Summary of genetic data and samples used in this and previous studies.

| Paper | Number of samples | Sampling locations | Genetic data |
| --- | --- | --- | --- |
| Warren et al. (2008) | 90 | QLD, NSW, VIC, TAS, South Australia | 57 retrotransposons |
| Kolomyjec et al. (2009) | 120 | 5 river systems in NSW, predominantly Hawkesbury and Shoalhaven | 12 microsatellites |
| Gongora et al. (2012) | 284 | 22 river systems across whole platypus range | haplotypes of mitochondrial control region and cytochrome b gene |
| Furlan et al. (2013) | 752 | 33 river systems across NSW, Victoria, TAS | 3 microsatellites, two mitochondrial haplotypes |
| this study | 57 | 12 river systems across QLD, NSW, TAS | 9.96 million SNPs from WGS data |

Using only a small number of markers gives limited information about the underlying population history of a species, and, because genealogies are stochastic given a particular demographic model, a genealogy built from a single locus, such as the mitochondrial control region (Gongora et al., 2012), may not reflect the historical relationships between populations under study (Novembre and Ramachandran, 2011). We anticipated that more could be learned about population structure and dynamics from whole-genome sequencing data, which contains considerably more information than the maternally-inherited mitochondria or than a few highly mutable microsatellites.

We sequenced the genomes of 57 platypuses from Queensland (QLD), New South Wales (NSW) and Tasmania (TAS) (Figure 1; Tables S1 and S2), in order to gain insights into the population genetics of the species. We investigated the differentiation between subpopulations, the relative historical population sizes and structure, and the extent of relatedness between the individuals sampled, which could be informative about the extent of individual platypus dispersal.

## Results

### Genome reassembly and SNP calling

We sequenced 57 platypus samples at 12-21X coverage, one in duplicate. We used the improved genome assembly ornAna2 (which will be made available no later than the time of publication of this paper) for all analyses, and ran standard software to jointly call variants across all samples (PLATYPUS (Rimmer et al., 2014)). The variant calls were filtered to produce a set of 6.7M stringently filtered SNPs across 54 autosomal scaffolds comprising 965Mb of the assembly.

### Data Quality

We undertook two different approaches to assess the quality of our SNP callset. In the first approach, two separate DNA samples from a single individual were sequenced. These were processed in identical fashion to all the other sequence data, with the processing blind to the fact that they were duplicates. After processing and SNP calling, we then compared the genotypes in the two samples from this individual. The rate of discordant genotypes between the two duplicate samples was very low (2.20 × 10^−3^ per SNP; 1.62 × 10^−5^ per bp; Table S3); because an error in either duplicate could lead to discordant genotypes, this would lead to an estimated error rate of 1.10 × 10^−3^ per SNP and 8.1 × 10^−6^ per bp. During our analyses we also discovered we had sampled a family quartet of two parents and two offspring (Table S4). This allowed us to use a second approach to assess the quality of our SNP callset by using the rate of Mendelian errors. We found a Mendelian error rate of 1.10 × 10^−3^ per SNP. In some configurations, an error in any one of the four genotypes would result in a Mendelian error; in others, an error would not be detected. Both approaches suggest an error rate of order 0.001 per genotype, suggesting that the dataset used in the analyses is of high quality. Our genotype error rate compares favourably with previous work in mountain gorillas (Xue et al., 2015) despite our lower sequencing coverage.

Sequencing a quartet also allowed us to estimate the switch error rate of haplotypes inferred in the new reference genome compared to the original assembly. Switch errors are changes in the pattern of inheritance in the two offspring, either due to errors or real recombination events (Browning and Browning, 2011). We found that the switch error rate was reduced by nearly 80% for ornAna2 compared to ornAna1 (Section S1; Table S5), indicating the new assembly contains far fewer errors. We conclude that the improved reference genome and stringent quality filters we have used mean that our dataset is of high quality.

### Relatives

Unlike earlier studies, our genome-scale data allowed us to look for relatives amongst our samples. Using the KING algorithm (Manichaikul et al., 2010) as implemented in VCFtools (Danecek et al., 2011), we identified many pairs of relatives (Table S4). In addition to a first-degree relative pair we had intentionally sequenced (a father-daughter pair from Taronga Zoo), we found we had sampled the aforementioned quartet from the Shoalhaven River, as well as the quartet mother’s sister. Additionally, there were 26 pairs of second- or third-degree relatives, in all cases from the same river or creek, or closely connected waterways, involving 28 of our 57 samples. For the analyses in this paper, except where otherwise noted, we used a set of 43 unrelated samples, indicated in Table S1.

### *De novo* mutation rate

We identified putative de novo mutations in the two offspring in the family quartet using the Bayesian filter incorporated into PLATYPUS, and further filtered them to remove any putative de novo mutation which was seen in any other sample. This gave us a total of 12 *de novo* mutations in the quartet, 6 in each offspring, and we estimated the *de novo* mutation rate at 7×10^−9^/bp/generation (95% CI 4.1×10^−9^−1.2×10^−8^/bp/generation). See Section S3 for more details.

### Dispersion and inbreeding

Of the quartet individuals, the mother, her sister and her two offspring were sampled in a pool at a junction about 2km downstream of the point where the father was found, which was in Jerrabattgulla Creek, a small tributary of the Shoalhaven River. Although the male offspring was first captured as a juvenile, because of difficulties in determining the age of adult platypuses, it is not possible to tell whether the two offspring were born as dizygotic twins or at different times, which would indicate that their parents mated in more than one year.

Given that we found so many instances of close relative pairs within the same river, it was natural to ask whether inbreeding is common in platypus. We examined long runs of homozygosity (LROHs) to investigate levels of inbreeding in the samples. Figure S1 shows *F_ROH_*, the estimated fraction of the analysed genome that is in LROHs (see Methods). The Carnarvon sample stands out, with *F_ROH_* estimated at 24.4%, but several other samples have *F_ROH_* higher than 10% (N745 and N711 from NQLD, N730 from the Gwydir River, N746 from the Broken River, N724 from the Barnard River, and N710 from Tasmania).

The length of homozygous segments depends on the recombination rate and number of generations since the most recent common ancestor of the two haplotypes. Since we do not know the fine-scale recombination rate in platypus, and estimation of segment length is complicated by the fragmented nature of the assembly, which may lead to ROHs being truncated artificially e.g. by scaffold ends (Figure S2), interpretation of the observed distribution of segment lengths is challenging. However, Figure S3 shows that the samples clearly fall into two groups: the north QLD (NQLD), central QLD (CQLD) and Gwydir samples (group 1) all have more ROHs than the other NSW and Tasmanian samples (group 2), but these have lower mean length than in some of the group 2 samples, and both groups contain samples with high overall *F_ROH_* This undoubtedly reflects differences in demographic history. In the case of the Carnarvon sample (N753), it is difficult to disentangle true inbreeding from low historical *N_e_*, since we only have one sample from this location, but the overall *F_ROH_* could be consistent with a mating between first-degree relatives. However, since N724 appears to be an outlier amongst the Barnard River samples, it may be that this individual is derived from a mating of individuals as closely related as second-degree. Thus, we cannot rule out the possibility of close inbreeding in wild platypus populations.

### Population structure

We first ran a Principal Component Analysis (PCA) to summarise the genetic variation, using the stringent SNP set after filtering based on minor allele frequency and missingness. Figure 2 shows the first two principal components (PCs). The first PC separates the Tasmanian from the mainland samples and accounts for 41.6% of the variation, and the second separates the mainland samples on a north-south axis and accounts for 22.2% of the variation. If we prune the SNPs based on linkage disequilibrium (LD), the first two PCs are exchanged and account for 27.6% and 23.7% of the variation (data not shown).

**Figure 2:**
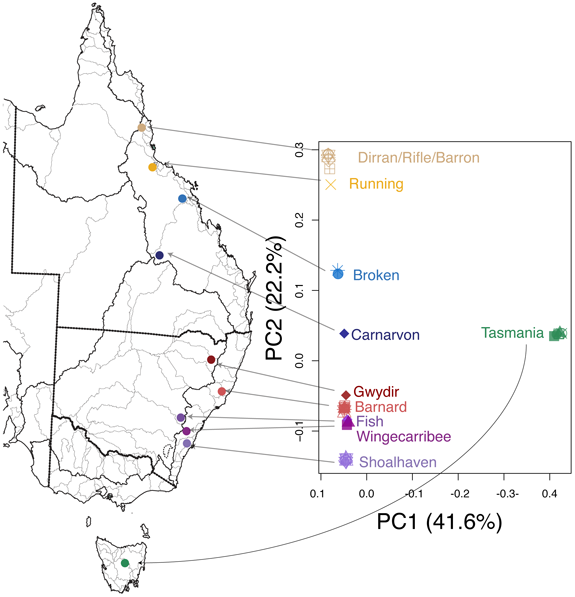
Principal component analysis on 43 unrelated samples. The first two principal components (PC1 and PC2) are shown, with the proportion of variance accounted for indicated in parentheses. The sampling locations are labelled in the same colour as the corresponding points. Samples from the same sampling location typically have extremely similar values on the first two PCs so that the corresponding points are often overplotted in the figure.

In order to explore differences between regions, we divided the samples into groups based on the PCA results and the samples’ known geographical proximity to one another (see Table S1). The Barnard River group has 11 individuals, and is combined with Gwydir River individual to form the north NSW group. The Shoalhaven River group has 12 individuals, and is combined with 2 individuals from the Wingecarribee River and one individual from the Fish River to form the central NSW group. Similarly, we grouped samples within each of Tasmania (N = 5) and NQLD (N = 7). The 3 samples from central QLD form their own group based on PCA and geography, and have been excluded from the following analyses due to low sample numbers.

The groups show different levels of nucleotide diversity, π (Nei and Li, 1979) (average number of nucleotide differences between individuals per site), ranging from 4.73 × 10^−4^ in the north QLD samples to about 1.02 × 10^−3^ in central NSW (CNSW) (Table 2). A large proportion of the SNPs segregating in each region are only polymorphic in that region (Figure S4): 34.5% of those in north QLD, 33.5% in north NSW (NNSW), 37.1% in central NSW and 72.1% in Tasmania (after downsampling to consider the same number of samples per region).

**Table 2:**
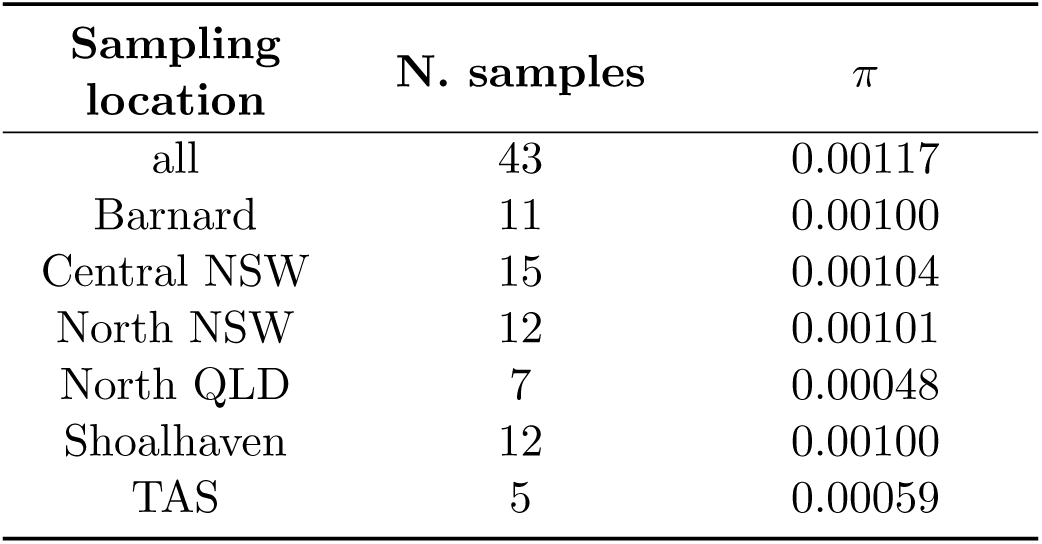
Nucleotide diversity (π) across different sampling locations. Note that North NSW is Barnard + Gwydir, and Central NSW is Shoalhaven + Wingecarribee + Fish Rivers. Central QLD has only 3 samples and is excluded from this analysis.

There was high *F_ST_* between the different regional groupings, with the highest value between Tasmania and north QLD (0.677) and the lowest between the Wingecarribee and Barnard groupings (0.077) (Table 3). The *F_ST_* values were slightly higher when we did not prune the SNPs based on local LD, since this unpruned SNP set retained many fixed differences between sampling locations (Table 3). There were a large number of fixed differences between both the Tasmanian and north QLD samples and the reference individual (Figure S5): 10% of the ~6.7 million SNPs segregating in the fifty-seven samples were fixed for the alternate allele in the five unrelated Tasmanian samples, and 7.3% were fixed in five randomly sampled unrelated north QLD samples.

**Table 3:** *F_st_* across different sampling locations. The black numbers above the diagonal are calculated using SNPs before LD pruning, and the blue ones below the diagonal are calculated after LD pruning (see Methods).

|  | Wingecarribee | North QLD | TAS | Barnard | Shoalhaven |
| --- | --- | --- | --- | --- | --- |
| Wingecarribee | - | 0.364 | 0.544 | 0.086 | 0.088 |
| North QLD | 0.335 | - | 0.726 | 0.362 | 0.394 |
| TAS | 0.445 | 0.676 | - | 0.555 | 0.555 |
| Barnard | 0.078 | 0.335 | 0.459 | - | 0.145 |
| Shoalhaven | 0.078 | 0.361 | 0.463 | 0.126 | - |

We applied STRUCTURE (Pritchard et al., 2000), a Bayesian model-based clustering algorithm, to identify subgroupings within our sampling location groups and assign the samples to them without using any prior information. Figure 3 shows the results from the admixture model in STRUCTURE, in which individuals are allowed to have membership in more than one of the *K* subgroupings, or clusters. We emphasise that inference of membership in multiple clusters does not necessarily mean that an individual has recent admixture; rather, these clusters represent putative ancestral populations which may have contributed to modern-day populations.

**Figure 3:**
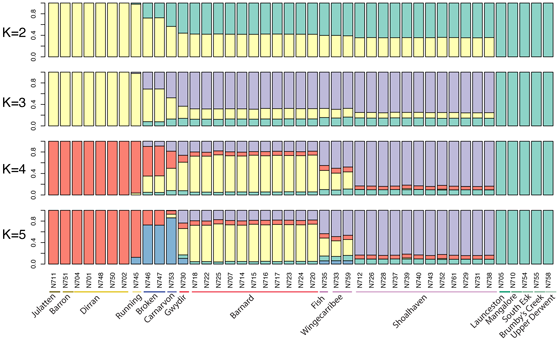
Population structure inferred from 43 unrelated individuals using STRUCTURE. Each individual is represented by a vertical bar partitioned into *K* coloured segments that represent the individual’s estimated membership fractions in *K* clusters. Ten STRUCTURE runs at each *K* produced very similar results, and so the run with the highest likelihood is shown. The Broken River and Carnarvon samples are labeled as central QLD and the Running River sample as north QLD, even though these samples did not form part of large clusters on the PCA and were excluded from the groupings used in Tables 2 and 3 and Figures S4 and S5.

For *K* = 2, the clusters are anchored by north QLD and Tasmania, with the central QLD and NSW groups inferred to be a mixture of these two. The next cluster at *K* = 3 corresponds to these central QLD and NSW individuals, and at *K* = 4, the north NSW (Barnard and Gwydir) individuals are delineated, along with individuals from the Fish and Wingecarribee Rivers. The additional cluster at *K* = 5 contributes the majority of the ancestry of the Carnarvon and Broken River samples, as well as a small amount of the ancestry of the samples from the Running River (from the Burdekin river system, like the Broken River individuals), and from the NSW rivers. When we ran STRUCTURE with *K* > 5, we found that the proportion of ancestry assigned to the additional clusters was always very low in all samples, and the posterior mean was close to 0 (i.e. the clusters were essentially empty), so, although slightly higher log likelihoods were observed for some runs at *K* = 8, we think that the simpler model with *K* = 5 represents the data better.

The most striking point in this figure is that the Wingecarribee and Fish River samples look more similar to the Gwydir and Barnard samples than they do to the Shoalhaven samples, despite being geographically adjacent to Shoalhaven (see Figure 1). On our PCA analyses, these samples fall in a position between the Shoalhaven and Barnard samples, but closer to the Barnard samples (rather than closer to Shoalhaven, as might be expected by their geographical location). We discuss possible explanations for the observed similarities below.

We used FineSTRUCTURE (Lawson et al., 2012) to further investigate population structure and demographic history. This method uses haplotype structure inferred from densely-typed markers to infer clusters of individuals with similar patterns of ancestry, and has been shown to be a particularly powerful approach to detecting fine-scale population structure (Leslie et al., 2015). A description of the FineSTRUCTURE algorithm can be found in the Methods section. Briefly, for each individual, the method first finds the other individuals who share ancestry most closely with it across different regions of the genome. Then, for each individual, the counts of the number of regions sharing most-recent-ancestry with each other individual form the rows of what is called a co-ancestry matrix. This information is then used to produce clusters of individuals with similar patterns of ancestry. Figure 4 shows the coancestry matrix and tree inferred by running FineSTRUCTURE on the 43 unrelated samples. The block structure of the coancestry matrix shows that there is strong population structure that is consistent with the samples’ geographic locations, but also shows evidence of finer-scale population structure within each river system.

**Figure 4.**
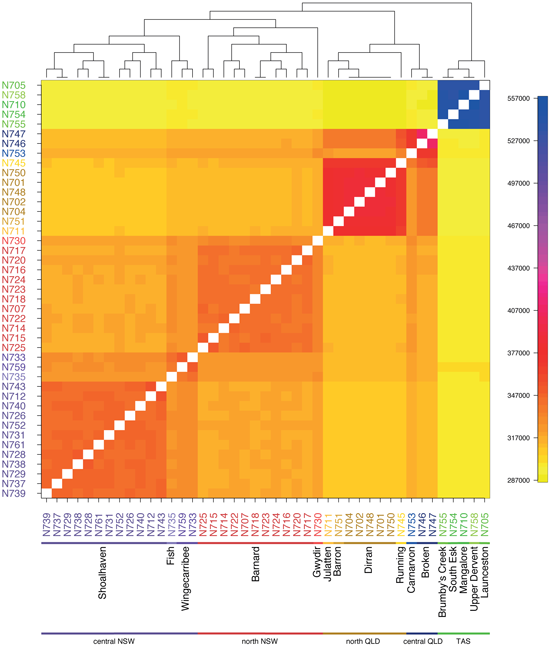
Coancestry matrix from 43 unrelated individuals using FineSTRUCTURE. Each row represents one of the sampled individuals, with the colours along the row for a particular individual representing the number of pieces of their genome for which each other individual shares most recent common ancestry with them. The tree shows the clusters inferred by FineSTRUCTURE from the coancestry matrix. The groupings on the x-axis are as in Figure 3.

The deepest branch on the tree separates the Tasmanian samples from the mainland, and the coancestry matrix shows little evidence of the sharing of most recent common ancestors between Tasmania and the mainland, implying a largely distinct population history for the Tasmanian samples, at least over the timescales during which they share recent ancestry with each other. The next branching splits the mainland samples into Queensland and New South Wales clusters, with a further split separating central NSW (including the Shoalhaven, Wingecarribee and Fish River samples) and north NSW (including the Gwydir River and Barnard samples). However, the Wingecarribee and Fish River samples show more haplotype sharing with the north NSW samples than with the Shoalhaven samples, supporting the evidence from STRUCTURE and PCA that these samples fall between the two larger clusters.

Fine-scale population structure is also evident within some river systems, with the samples from Shoalhaven and Barnard rivers subdivided into smaller population clusters. By contrast, the five samples from the Dirran River in north QLD form a single cluster. The single sample from the Carnarvon River (N753), while showing the greatest level of haplotype sharing with the other central QLD sample from the Broken River, also shows more sharing with the samples from NSW and less with the samples from north QLD than would be expected based on geography, as they are closer to the north QLD rivers than those in NSW. By contrast, the Broken River samples show much greater sharing with the north QLD samples than the Barnard River samples, as expected. We hypothesize that this may be due to ancestral admixture between the Broken and Carnarvon rivers, and subsequent admixture between the north QLD samples and the Broken River samples only.

We further investigated demographic history using the Pairwise Sequentially Markovian Coalescent (PSMC) method of Li and Durbin (2011). PSMC examines how the local density of heterozygous sites changes along the genome, reflecting chromosomal segments of constant time to the most recent common ancestor (*T_MRCA_*), separated by recombination events. Knowing the coalescence rate in a particular epoch allows estimation of *N_e_* at that time. As has become clear from applications in other contexts (e.g. Prado-Martinez et al., 2013; Wallberg et al., 2014; Nadachowska-Brzyska et al., 2016), this comparison of the two chromosomes within a diploid genome offers an extremely powerful tool for inferring historical effective population size. The power of this approach lies in the fact that there are many thousands of “replicate” segments within a single diploid genome, and these collectively provide precise estimates of historical population size, except for the very recent past and the distant past.

Figure 5 shows estimates of the effective population size, *N_e_*, from each sample at a series of time intervals. The scaling on the X-axis of this plot depends on the generation time, *g*, and the scaling of both the X- and Y-axes depend on the mutation rate *μ*. Little is known about these two parameters for the platypus, but changing them will affect the estimates for all samples equally, by simply linearly rescaling the axes (Figures S6 and S7). We will thus focus primarily on conclusions based on relative differences between PSMC estimates as these do not depend on assumptions about *g* and *μ*. For scaling the axes in Figure 5, we used *g* = 10 years, following Furlan et al. (2012), which is consistent with the known observations that platypus can live up to 20 years in the wild and that both sexes can reproduce from the age of 2 years, although first breeding in some females can be later than this age (Grant, 2004a, 2007). In Figure 5, we used a mutation rate of 7 × 10^−9^/bp/year, the *de novo* mutation rate we estimated from our own data using the quartet. Note that, in the time-scaling used in Figure 5, the method is not informative more recently than about 10,000 years ago, or further into the past than about 1-2M years.

**Figure 5:**
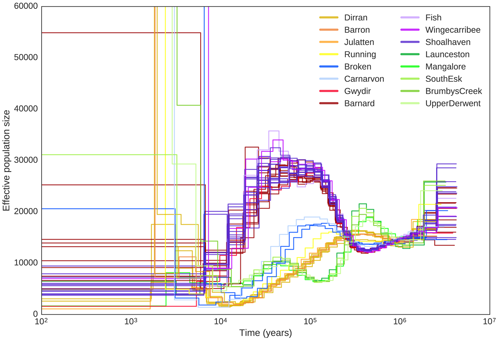
Historical effective population sizes inferred using PSMC. Each line represents a single individual with lines coloured according to sampling location. Trajectories were scaled using *g* = 10 and *μ* = 7 × 10^−9^. Effective population size was truncated at 60,000. Samples from a similar sampling location show very similar trajectories.

Individuals from the same river show strikingly similar trajectories in Figure 5, supporting the precision of the relative *N_e_* estimates and giving us confidence that we are measuring real features of population history. Bootstrapping performed according to the method in Li and Durbin (2011) shows similar trajectories over 100 replicates for each sample (Figure S8) with the exception of very recent and very distant time points (more recent than 5-10,000 years and older than 1M years), as expected (Li and Durbin, 2011). The highly congruent trajectories within a river system suggest that, looking backwards in time, the ancestors of these samples were probably part of the same population within the timeframe accessible to the method. Although samples from the same population would be expected to show the same *N_e_* trajectory, having the same *N_e_* trajectory does not necessarily mean that samples are from the same population. However, having different *N_e_* trajectories around a certain time in the past is difficult to reconcile with the individuals’ ancestors coming from the same ancestral population at that time.

One striking feature of Figure 5 is that there are clearly four distinct groups of samples (all NSW, central QLD, north QLD, and Tasmania, respectively) with the ancestors of each group clearly having separate population histories until well into the past. It is around 1M years (in the time-scaling of Figure 5) before NSW, north QLD, and Tasmania begin to share ancestral history, and perhaps 300,000 years until the central QLD and NSW samples might share ancestral history. This implies that there has been extensive population structure in platypus samples across Australia over a long time period.

A second feature of Figure 5 is that all *N_e_* trajectories take their lowest values at the most recent time points for which the method is informative. This would be consistent with a decline in platypus numbers across Australia over the time period accessible to the method. In the case of the north QLD samples, the *N_e_* level becomes extremely low and remains so, and corresponds to a marked population bottleneck (around 10,000 years ago in this time-scaling). This is consistent with the low nucleotide diversity observed in these populations, and with these samples clustering as a single homogenous population in the FineSTRUCTURE results. The central QLD population may well also have been affected by a recent bottleneck and shows very low *N_e_* during this period.

## Discussion

We have described the first population-scale whole-genome sequencing study of the platypus. The analyses presented here provide insights into the population structure and levels of diversity in this species not previously possible with microsatellite markers or mtDNA. Table 1 provides details of the source and extent of genetic variation used in this and previous studies of platypus demography and population structure.

Our whole-genome data allowed us to estimate relatedness between individuals, and we found that more than half of our samples had a least a third-degree relative amongst the other individuals sampled from the same river. The quartet samples were all collected within a small distance of each other over a relatively short timeframe (< 3 years). These observations are consistent with an underlying pattern of limited dispersal of (at least some) relatives. This is somewhat surprising given that previous studies using mark-recapture approaches in the Shoalhaven River (Grant, 2004b; Bino and Grant, 2015) and other streams (Serena, 2012; Serena et al., 2014) have reported the dispersal of a high proportion of juveniles, especially males, few recaptures of adult males, and the continued capture of unmarked males and females. Of all the pairs of relatives we sampled at the same site, most were female-female or male-female, and not male-male (there are only three male-male relative pairs in Table S4), which is consistent with male-biased dispersal.

Since it appears to be common to collect related individuals when sampling at the same location across several years, it is likely that previous population genetic studies on the platypus (Kolomyjec et al., 2009; Gongora et al., 2012; Furlan et al., 2013) included relatives, but that this was not detected due to the small number of markers used. Some of these studies included individuals in our sequencing study. It is unclear whether and to what extent the results from these earlier studies, particularly those from model-based approaches, would have been affected by the inadvertent inclusion of close relatives.

The inclusion of a quartet in our samples allowed us to estimate the *de novo* mutation rate in the platypus, the first estimate in a non-placental mammal. While our estimates were limited to only a single quartet sequenced to moderate coverage, our estimate is consistent with previous work to estimate mutation rates in mammals. Our point estimate of the rate of 7.0 × 10^−9^ (95% CI 4.1×10^−9^−1.2×10^−8^/bp/generation) is lower than the estimated rate of 1.2 × 10^−8^ in humans and chimpanzees (Kong et al., 2012; Venn et al., 2014) but higher than the rates estimated for laboratory mice (5.4 × 10^−9^) (Uchimura et al., 2015). The relative ordering of the point estimates is consistent with the observation that mutation rates in mammals are negatively correlated with body mass and generation time (Welch et al., 2008), but given the various sources of uncertainty in our estimate (some of which are not easy to quantify) it would not be appropriate to place undue weight on it.

For population genetic analyses, descriptive approaches have some advantages over those based on population genetic models. Where descriptive approaches point to clear conclusions, there can be confidence that these are valid. Methods based on population genetics modelling can be more powerful, but their interpretation is always complicated by the fact that they depend, often to a degree which is hard to assess, on the underlying model assumptions. We have thus focused primarily on approaches which do not rely on population genetics modelling. (Some of the approaches used above, e.g. STRUCTURE, and FineSTRUCTURE, do involve statistical models which aim to capture features of the data, but none is based on models of historical population dynamics or history.) Our analyses confirm the strong structure previously reported in the species (Gongora et al., 2012; Furlan et al., 2013), with both pronounced differentiation on the mainland in a north-south direction, and separation of the Tasmanian samples from all other groups. Consistent with Furlan et al. (2013), we saw little evidence for structure within the Tasmanian samples, which could be due to greater overland migration in this wetter climate. On the other hand, the apparent lack of structure could simply be because of our small sample size (6 individuals).

Our results suggest several instances of higher genetic similarity between individuals than expected given their sampling locations. For example, the Fish and Wingecarribee samples come from rivers that flow into different systems on either side of the Great Dividing Range (the west-flowing Murray-Darling system and the east-flowing Hawkesbury system), and yet look extremely similar to one another on the PCA and in STRUCTURE, and are grouped together in the FineSTRUCTURE analysis. This finding is consistent with Furlan et al. (2013), who also reported that there was little or no genetic differentiation in mtDNA and microsatellite markers between platypus either side of the Great Dividing Range in Victoria. This genetic similarity might be due to overland migration or historical connections between the Murray-Darling and Hawkesbury systems that have been altered due to climate or geological changes. The similarity between the sample from Rifle Creek, which is part of the west-flowing Mitchell River system, and those from east-flowing Dirran Creek and Barron River, could probably be explained by occasional overland dispersal. Although they are part of different drainages, the sampling locations are all within 100km, and the climate is much wetter, so migration may have been possible if the ephemeral streams and refuge pools in the area persisted for long enough. The similarity may also be due to the population decline evident from the PSMC results, and low genetic diversity across the north QLD samples as a whole.

It is interesting that Kolomyjec et al. (2009) reported that 13 of the 120 individuals they analysed at 12 microsatellites appeared to be first-generation migrants from the Shoalhaven to the Hawkesbury River systems (the latter including Wingecarribee), or vice versa. (Where sampling includes close relatives, as seems possible in the light of our results, estimating numbers of migrants may not be straightforward.) We did not find any evidence of migrants nor of recent admixture (using the PCA or STRUCTURE) between the Shoalhaven and the Wingecarribee, consistent with the different river systems being separated by steep terrain along much of their border, despite the physically close sampling locations. It may be that we simply did not happen to sample such individuals, or that the sampling locations were further apart than those of Kolomyjec et al. (2009).

The PSMC results provide a glimpse of the past demographic histories of the different sampling locations, which was not possible in previous studies based on a small number of markers. The fact that these groups show such different histories emphasises that running this method on a single sample from a species might not provide a representative picture of the coalescence process for that species, as noted by Nadachowska-Brzyska et al. (2016) in a study on flycatchers. Our PSMC results reveal a recent reduction in *N_e_* in all regions, with a particularly low *N_e_* in the Queensland samples.

The Queensland bottleneck likely reflects the historical and current isolation and paucity of suitable habitat for platypus between North (Australian Wet Tropics) and Central QLD, known as the “Burdekin gap” (named for the Burdekin River). This hot and dry area is currently climatically unsuitable for platypus (Figure S14) and has long acted as a barrier to genetic exchange (James and Moritz, 2000; Schauble and Moritz, 2001). The declining *N_e_* but separate trajectories of the north and south QLD samples reflect accumulating evidence of the impact of arid periods - glacial maxima - on the mesic forest biotas of the region (Bryant and Krosch, 2016). South of this, the Broken River is within the mid-east QLD diversity hotspot for rainforest faunas, and the nearby coast has functioned as a small and isolated climate change refugium since the Last Glacial Maximum. Carnarvon Gorge is an oasis in the semi-arid heart of central QLD, making it the only suitable current habitat for platypus for hundreds of kilometres (Figure S14). The possible ancestral admixture between the Carnarvon and Broken river systems, and subsequently between the Broken River and other north QLD samples, is hard to reconcile with the current climate of these areas. The high level of homozygosity seen in the Carnarvon sample may reflect the low effective population size over the last ~50,000 years (Figure 5) or may be the result of recent inbreeding. Regardless, it suggests that the Carnarvon River platypuses may be particularly vulnerable and should therefore be a priority for conservation, as noted by Kolomyjec et al. (2013).

In contrast to the QLD samples, the NSW individuals appear to have had higher and relatively stable *N_e_* over much of their history, with population decline only in more recent time periods. The high *N_e_* makes sense given recent paleoecological evidence that parts of this region remained wetter (relative to North QLD and Southeast Australia) during the Last Glacial Maximum (Moss et al., 2012) and, based on paleoclimate modeling, the region is inferred to have been a large, stable mesic refuge over the past 120,000 years (Weber et al., 2014; Rosauer et al., 2015). Determining whether the recent decline in *N_e_* is due to late Pleistocene climate change or other causes would require high resolution modeling of paleoclimates for this region.

PSMC results can be used to infer when different populations separated by noting where the *N_e_* estimates start to diverge when moving forward in time (right to left in Figure 5) (Li and Durbin, 2011; Nadachowska-Brzyska et al., 2013; Thomas et al., 2015). Regardless of the scaling of the axes, Figure 5 implies that all of the NSW samples share an ancestral population more recently than any of them shares an ancestral population with any of the other samples. Looking backwards in time, the central QLD samples next share an ancestral population with the NSW samples, before this larger group share an ancestral population with either the north QLD or Tasmanian samples. There are slight differences in *N_e_* trajectories which persist throughout the timescales covered by Figure 5. (As noted above, we would expect, and see, differences at the extreme right of the figure reflecting noise associated with the loss of precision of the estimates in the ancient past.) Between about 1 and 2 million years ago in the time-scaling of the figure, the *N_e_* trajectories of each of the Tasmanian, North QLD, NSW, and Central QLD samples are extremely close, but not identical. Further, the differences are typically larger between than within these three groupings. One explanation for this is that all groupings do share an ancestral population by this time, but that there are systematic differences between the estimates from each which result in the small differences in *N_e_* trajectories between groups. The other explanation is that the groupings do not share an ancestral population at that time, but instead their ancestral populations just happened to have extremely similar population sizes. This second explanation would be consistent with the divergence of *N_e_* estimates on the right of Figure 5, where the values for the north QLD and Tasmanian samples jump to higher values (looking left to right) before those from NSW/central QLD. In our view, limited weight should be put on this last observation because *N_e_* estimates are noisy at the right of the picture, and the nature of this noise could differ systematically between groupings for many reasons, including systematic differences in properties of SNPs in those groups. Our data cannot distinguish between these two explanations (sharing of an ancestral population by all samples, or distinct ancestral populations with extremely similar *N_e_* values over long time periods), but the former explanation is more parsimonious and appears to us to be the more likely.

Regardless of the preferred explanation of the two in the previous paragraph, there is no evidence in our data that the north QLD, central QLD, and NSW samples shared an ancestral population before either shared an ancestral population with the Tasmanian samples. Under the first explanation, all three groupings first shared an ancestral population around the same time. Under the second explanation, none of them share an ancestral population over the time scales for which reliable PSMC *N_e_* estimates are available. We think it is most likely that there were three ancestral populations (TAS, north QLD and north NSW/central QLD) which all coalesced around the same time (about 800KYA, using the scaling in Figure 5). This is hard to reconcile with the phylogenetic tree inferred by Gongora et al. (2012) using mitochondrial data, in which all north and central QLD samples (including from Dirran Creek, Running River and Carnarvon) coalesce much more recently with each other than they do with any other population. On the other hand, the wide 95% credible intervals for divergence times inferred by Gongora et al. (2012) overlap between the different events, and our estimates from Figure 5 fall within them. Although mitochondrial ancestries could indeed differ from population ancestries and those of autosomal loci, the estimates from PSMC are expected to be more precise than those available from any single locus, since the PSMC method, in effect, averages over every locus in the genome.

Interestingly, the divergence times we and Gongora et al. (2012) have estimated predate the earliest fossil evidence for platypus (Musser, 1998, 2013), although we are conscious that our absolute estimates do depend on the generation time and mutation rate. This finding does not necessarily contradict fossil evidence but suggest that the modern platypus extends back to the Early to Middle Pliocene. This could be consistent with it having evolved from the giant platypus species *O. tharalkooschild*, as suggested by Pian et al. (2013).

We have presented the first whole-genome resequencing study of the platypus. Our results have provided insights into the strong population structure in this species as well as the demographic history of the different sampling locations, at a level that was impossible with older molecular tools. Future studies are likely to shed further light on the population history and biology of this fascinating species. This study emphasises the power of whole genome sequencing for inference of population dynamics and historical demography, even where sampling of individuals is constrained.

## Methods

### Samples and sequencing

We obtained DNA samples from 61 platypuses, extracted from toe webbing, spleen, liver or cell lines. These included between one and eighteen individuals from each of seventeen waterways across most of the platypus range, excluding Victoria because no high-quality DNA samples were available. For quality control, we included a duplicate sample from Dirran Creek, north QLD, and a father-daughter pair obtained from Taronga Zoo (the father and mother originally being from the Fish River in the Macquarie Basin in NSW). These were sequenced in four tranches. Details of the library preparation and paired-end (PE) sequencing are shown in Table S6. Three samples were discarded because they failed sequencing quality control. Mean coverage and insert size for the remaining 58 samples are shown in Figure S15 and Table S2.

### Reference genome reassembly

All analyses in this paper were carried out on the platypus male reference genome assembly, ornAna2. The new platypus reference genome will be made available no later than the time of publication of this paper.

### Mapping and variant calling

We mapped the paired-end data from all 58 samples to ornAna2 using Stampy (Lunter and Goodson, 2011) without BWA pre-mapping. Duplicates were removed using Picard MarkDuplicates tool (http://broadinstitute.github.io/picard). Variant calling was performed jointly on the 58 samples using the PLATYPUS variant caller (Rimmer et al., 2014), and we obtained a total of 14,127,611 biallelic SNPs with the “PASS” filter. We removed indel calls, monomorphic positions, and 2042 positions where the reference individual (N720) was called homozygous for the alternative allele, which may represent errors in the reference genome. Inspection of the variant calls which were discordant between duplicate samples (see section S2) revealed that PLATYPUS incorrectly called positions as heterozygous despite no reads supporting the variant at that position. We thus removed 223,103 variants across all individuals studied for which PLATYPUS reported no reads supporting the variant. We assessed sites for Hardy-Weinberg Equilibrium, as implemented in VCFtools (0.1.14), and excluded SNPs with p-value below 10^−7^. These variants may be errors but the substantial population structure in our samples may also cause variants to fail this test. We filtered out variants with quality below 60, and kept only SNPs with no missing data. This SNP set forms the stringent SNP set referenced in the results.

To assign contigs to chromosomes, we took advantage of the chromosome assignment made for the ornAna1 reference, in which sequences where attributed to chromosomes 1 to 7, 10-12, 14, 15, 17, 18, 20, X1, X2, X3 and X5. Specifically, we broke down each chromosome from ornAna1 in pieces of 500Kb and aligned these pieces to the new reference genome using bwa mem (BWA version 0.7.12). We excluded the pieces that resulted in primary and secondary alignment, because these are likely to be due to mis-assemblies in ornAna1. For the remaining pieces, we kept only the primary alignment with mapping quality of 60 (highest value). A total of 304 contigs, with total sequence length of 1,779,183,769 bp, had at least one piece of ornAna1 chromosome aligning to them with these criteria. The remaining 4268 contigs (211,276,236 bp) were excluded from further chromosome assignment using homology to ornAna1. For each contig, we computed the number of base pairs covered by pieces coming from each of the ornAna1 chromosomes. We assigned the ornAna1 chromosome label to a contig when more than 90% of covered sequence within the contig was attributed to the chromosome in ornAna1 and when more than 10% of the total sequence of the contig was covered by ornAna1 sequence pieces. Finally, we compared the coverage for each contig between males and females to validate the autosomal contigs. For subsequent analyses, we retained only 54 assigned autosomal contigs with at least 50 SNPs that passed our filters, covering 965,354,475 bp in total. This final SNP callset includes a total of 6,727,617 SNPs.

We determined the callable regions of the genome by running PLATYPUS with the callRefBlocks option to determine the bases where a good quality reference call could be made, and combined these with the SNP callset positions to give a callable genome length of 910,563,690bp.

### Identifying relatives

To identify relatives among the sampled individuals, we ran the KING algorithm (Manichaikul et al., 2010) as implemented in VCFtools (0.1.14) (Danecek et al., 2011), which uses the number of SNPs that are identical-by-state 0, 1 or 2 to estimate the kinship coefficient. For each broad sampling location (north QLD, north NSW, central NSW, and Tasmania) we removed SNPs with minor allele frequency (MAF) < 0.05 across samples from that location. For several of the population genetic analyses described below (*F_ST_* and nucleotide diversity calculations; STRUCTURE; FineSTRUCTURE), we removed one individual from each relative pair (up to and including third-degree relatives), leaving 43 individuals (shown in blue in Table 4.2). Specifically, we removed N713, N719, and N721 from the Barnard River; N703 and its duplicate, N749, from Dirran Creek; N727, N734, N741, N742, N757 and N760 from the Shoalhaven River; N732 and N744 from the Wingecarribee River; N736, the daughter from the Fish River (Taronga Zoo); and N756 from Brumby’s Lake in Tasmania.

### Long runs of homozygosity

We followed the approach of Prado-Martinez et al. (2013) to detect long regions of homozygosity (LROHs). For each sample, we calculated the heterozygosity (i.e. proportion of callable sites that were called as heterozygous) in overlapping windows along the genome, using the set of callable sites defined above. We examined the distribution across windows, selected a threshold (we chose 5 × 10^−5^, based on the local minima in Figure S16) below which a window was classed as “homozygous”, then merged overlapping homozygous windows. We used 1Mb windows, shifted by 200kb each time, and used only the scaffolds classed as autosomal that were longer than 1Mb (total length 963.8Mb). We then calculated the proportion of the examined genome that was in homozygous chunks, *F_ROH_*, as a measure of inbreeding (Figure S1).

### Population genetic analyses

#### Nucleotide diversity, *F_ST_* estimates, PCA and STRUCTURE

We used VCFtools (0.1.14) to compute π per site (`--site-pi`) on all SNPs in unrelated individuals from each sampling location. The total nucleotide diversity for a sampling location was computed by summing over values of π for all polymorphic SNPs and dividing by the total number of callable sites (910,563,690).

We used EIGENSOFT (v6.0.1) (Patterson et al., 2006) to estimate *F_ST_* between sampling locations and to run a Principal Components Analysis (PCA). We ran STRUCTURE (Pritchard et al., 2000) with the admixture model for *k* = 2, 3,…10. For these analyses, we took the callset on the unrelated samples and removed SNPs with MAF < 0.05 (leaving 3,245,503 SNPs), then carried out LD pruning using PLINK (Purcell et al., 2007). Specifically, for windows of 50 SNPs, we removed one of each pair of SNPs if the *r*^2^ between them was greater than 0.1, and then shifted the window 5 SNPs forward (PLINK option: `--indep-pairwise 50 5 0.1`). This left us with 128,803 SNPs.

#### Phasing and FineSTRUCTURE

We used SHAPEIT2 (O’Connell et al., 2014) to phase haplotypes across samples. SHAPEIT2 was run on the full cleaned SNP set (i.e. before MAF filtering and LD pruning) using a window size of 0.5Mb, 200 conditioning states, and 30 iterations of the main MCMC, and incorporating phase-informative sequencing reads to improve phasing at rare variants (Delaneau et al., 2013). The haplotypes were post-processed with duoHMM (O’Connell et al., 2014), using the two known pedigrees in the sample set to improve the phasing and correct Mendelian errors.

The phased haplotypes were used to run FineSTRUCTURE (Lawson et al., 2012). The FineSTRUCTURE algorithm involves two separate stages. The first stage considers each sampled individual separately, and proceeds along the phased chromosomes (or, in our case, scaffolds) in each individual. For a particular individual, the method partitions the chromosome into pieces, and for each such piece it searches amongst the other sampled individuals to find the one who shares most recent common ancestry for that part of the chromosome. (The partitioning of the phased chromosomes into pieces, and the identification of the piece in another individual sharing most recent common ancestry, are undertaken jointly in the algorithm.) For this individual, one can then count the number of chromosomal pieces for which each other individual shares most recent common ancestry. This process is undertaken separately for each sampled individual. These shared ancestry counts can be visualized in what is called a coancestry matrix. The second stage of the FineSTRUCTURE algorithm involves taking these coancestry counts for each individual, and forming clusters of individuals with the property that individuals within the same cluster have similar patterns of sharing with other individuals. One can then obtain a tree relating the sampled individuals by successively merging clusters which are similar. (When FineSTRUCTURE produces the tree, similarity of clusters is defined in terms of changes in the likelihood of the statistical model used by the algorithm to infer clusters. The tree should not be interpreted as a direct estimate of the ancestral history of the samples.)

FineSTRUCTURE was run using the linked model with a uniform recombination rate and default parameters except 2M iterations of the MCMC (1M for burn-in), 200,000 tree iterations, and starting *N_e_* of 10M to avoid the EM algorithm finding a local optima with zero recombination. Three MCMC runs were performed, giving identical sample clustering.

### PSMC

We ran PSMC to investigate historical population sizes (Li and Durbin, 2011). A diploid fasta file was created from the SNP set referenced above, and used to run PSMC for each sample. PSMC was run for 25 iterations using an initial *θ*/*ρ* ratio of 5 and the default time patterning. Bootstrapping was performed as in Li and Durbin (2011) by resampling 5Mb chunks of the genome with replacement to generate 30 artifical chromosomes of 100Mb each and running 100 bootstrap replicates.

### Habitat modelling

We used the maximum-entropy approach of Phillips et al. (2006) to model the platypus habitat (Figure S14). This method uses a set of layers, or environmental variables, as well as a set of geo-referenced occurrence locations, to produce a model of the range of a given species. We used the following variables: Precipitation - annual mean; Temperature - annual max mean; Temperature - annual min mean; Drainage - variability; Drainage - average; Drainage Divisions Level 1; Drainage Divisions Level 2; River Regions.

## Acknowledgements

We thank Weerachai Jaratlerdsiri for helping to prepare DNA samples, Freya Shearer (University of Oxford) for help with Figure S14, Russell Jones (University of Newcastle), Josh Griffiths and Nick Gust (Department of Primary Industries and Water Hobart, Resource Management and Conservation Division, Tasmania) for providing samples, Jo Wiszniewski (Taronga Zoo) for advice and Wes Warren (Washington University) for his advice on genome assembly. We are grateful to Guojie Zhang (University of Copenhagen) and the Genome 10K Consortium for their collaboration and making available data on the new platypus reference genome. We thank the High-Throughput Genomics Group at the Wellcome Centre for Human Genetics (funded by Wellcome Trust grant reference 090532/Z/09/Z) for the generation of sequencing data.

This work was supported by a Wellcome Trust Core Award (090532/Z/09/Z) to P.D. and by a University of Sydney Start-Up Research grant to J.G.

## Data Availability

Data will be released on publication of the paper.

## Author Contributionss

P.D. and J.G. conceived the study. P.W., T.D., S.K., M.S., T.G., F.G., and J.G. contributed samples and performed experiments. P.P. and R.B. performed library preparation and led the sequencing. H.C.M., E.M.B., and J.H. performed the analyses. H.C.M., E.M.B., J.H., P.D., C.M., and J.G. drafted the paper. P.D. supervised the project. All authors read and commented on the manuscript.

## References

Abascal, F., Corvelo, A., Cruz, F., Villanueva-Cañas, J. L., Vlasova, A., Marcet-Houben, M., Martínez-Cruz, B., Cheng, J. Y., Prieto, P., Quesada, V., Quilez, J., Li, G., García, F., Rubio-Camarillo, M., Frias, L., Ribeca, P., Capella-Gutiérrez, S., Rodríguez, J. M., Câmara, F., Lowy, E., Cozzuto, L., Erb, I., Tress, M. L., Rodriguez-Ales, J. L., Ruiz-Orera, J., Reverter, F., Casas-Marce, M., Soriano, L., Arango, J. R., Derdak, S., Galán, B., Blanc, J., Gut, M., Lorente-Galdos, B., Andrés-Nieto, M., López-Otín, C., Valencia, A., Gut, I., García, J. L., Guigó, R., Murphy, W. J., Ruiz-Herrera, A., Marques-Bonet, T., Roma, G., Notredame, C., Mailund, T., Albà, M. M., Gabaldón, T., Alioto, T., and Godoy, J. A. (2016). Extreme genomic erosion after recurrent demographic bottlenecks in the highly endangered iberian lynx. Genome Biology, 17(1): 251.

Bick, Y. and Sharman, G. (1975). The chromosomes of the platypus *(Ornithorhynchus*: Monotremata). Cytobios, 14:1728.

Bino, G. and Grant, T.R. & Kingsford, R. (2015). Life history and dynamics of a platypus (Ornithorhynchus anatinus) population: four decades of mark recapture surveys. Sci Rep, 5:16073.

Bosse, M., Megens, H.-J., Madsen, O., Frantz, L. A., Paudel, Y., Crooijmans, R. P., and Groenen, M. A. (2014). Untangling the hybrid nature of modern pig genomes: a mosaic derived from biogeographically distinct and highly divergent Sus scrofa populations. Molecular ecology, 23(16):4089–4102.

Browning, S. R. and Browning, B. L. (2011). Haplotype phasing: existing methods and new developments. Nat Rev Genet, 12(10):703–714.

Bryant, L. M. and Krosch, M. N. (2016). Lines in the land: a review of evidence for eastern australia’s major biogeographical barriers to closed forest taxa. Biological Journal of the Linnean Society, 119(2):238–264.

Danecek, P., Auton, A., Abecasis, G., Albers, C. A., Banks, E., DePristo, M. A., Handsaker, R. E., Lunter, G., Marth, G. T., Sherry, S. T., McVean, G., Durbin, R., and (2011). The variant call format and vcftools. Bioinformatics, 27(15):2156.

Delaneau, O., Howie, B., Cox, A., Zagury, J.-F., and Marchini, J. (2013). Haplotype Estimation Using Sequencing Reads. The American Journal of Human Genetics, 93(4):687–696.

Freedman, A. H., Gronau, I., Schweizer, R. M., Ortega-Del Vecchyo, D., Han, E., Silva, P. M., Galaverni, M., Fan, Z., Marx, P., Lorente-Galdos, B., Beale, H., Ramirez, O., Hormozdiari, F., Alkan, C., Vilà, C., Squire, K., Geffen, E., Kusak, J., Boyko, A. R., Parker, H. G., Lee, C., Tadigotla, V., Wilton, A., Siepel, A., Bustamante, C. D., Harkins, T. T., Nelson, S. F., Ostrander, E. A., Marques-Bonet, T., Wayne, R. K., and Novembre, J. (2014). Genome sequencing highlights the dynamic early history of dogs. PLoS Genet, 10(1):e1004016.

Furlan, E., Griffiths, J., Gust, N., Handasyde, K., Grant, T., Gruber, B., and Weeks, A. (2013). Dispersal patterns and population structuring among platypuses, Ornithorhynchus anatinus, throughout south-eastern Australia. Conservation Genetics, 14(4):837–853.

Furlan, E., Stoklosa, J., Griffiths, J., Gust, N., Ellis, R., Huggins, R. M., and Weeks, A. R. (2012). Small population size and extremely low levels of genetic diversity in island populations of the platypus, Ornithorhynchus anatinus. Ecol Evol, 2(4):844–857.

Gongora, J., Swan, A. B., Chong, A. Y., Ho, S. Y., Damayanti, C. S., Kolomyjec, S., Grant, T., Miller, E., Blair, D., Furlan, E., et al. (2012). Genetic structure and phylogeography of platypuses revealed by mitochondrial DNA. Journal of Zoology, 286(2):110–119.

Grant, T.R. G. M. T.-S. P. (2004a). Breeding in a free-ranging population of platypuses, Ornithorhynchus anatinus, in the upper Shoalhaven River in New South Wales - a 27 year study. Proceedings of the Linnean Society of NSW.

Grant, T. (2004b). Captures, capture mortality, age and sex ratios of platypuses,Ornithorhynchus anatinus, during studies over 30 years in the upper Shoalhaven River in New South Wales. Proceedings of the Linnean Society of NSW.

Grant, T. (2007). Platypus. Australian Natural History Series. CSIRO Publishing, 4th edition.

Grutzner, F., Rens, W., Tsend-Ayush, E., El-Mogharbel, N., O’Brien, P. C., Jones, R. C., Ferguson-Smith, M. A., and Marshall Graves, J. A. (2004). In the platypus a meiotic chain of ten sex chromosomes shares genes with the bird Z and mammal X chromosomes. Nature, 432(7019):913–7.

Grutzner, Frank Rens, Willem Tsend-Ayush, Enkhjargal El-Mogharbel, Nisrine O’Brien, Patricia C M Jones, Russell C Ferguson-Smith, Malcolm A Marshall Graves, Jennifer A England Nature. 2004 Dec 16;432(7019):913–7. Epub 2004 Oct 24.

Gust, N., Griffiths, J., Driessen, M., Philips, A., Stewart, N., and Geraghty, D. (2009). Distribution, prevalence and persistence of mucormycosis in tasmanian platypuses (ornithorhynchus anatinus). Australian Journal of Zoology, 57(4):245–254.

Hufford, M. B., Xu, X., Van Heerwaarden, J., Pyhäjärvi, T., Chia, J.-M., Cartwright, R. A., Elshire, R. J., Glaubitz, J. C., Guill, K. E., Kaeppler, S. M., et al. (2012). Comparative population genomics of maize domestication and improvement. Nature genetics, 44(7):808–811.

Isaac, N. J., Turvey, S. T., Collen, B., Waterman, C., and Baillie, J. E. (2007). Mammals on the edge: Conservation priorities based on threat and phylogeny. PLOS ONE, 2(3):1–7.

James, C. and Moritz, C. (2000). Intraspecific phylogeography in the sedge frog Litoria fallax (Hylidae) indicates pre-Pleistocene vicariance of an open forest species from eastern Australia. Molecular Ecology, 9(3):349–358.

Klamt, M., Thompson, R., and Davis, J. (2011). Early response of the platypus to climate warming. Global Change Biology, 17(10):3011–3018.

Kolomyjec, S., Grant, T., Johnson, C., and Blair, D. (2013). Regional population structuring and conservation units in the platypus (Ornithorhynchus anatinus). Australian Journal of Zoology, 61:378385.

Kolomyjec, S. H., Chong, J. Y., Blair, D., Gongora, J., Grant, T. R., Johnson, C. N., and Moran, C. (2009). Population genetics of the platypus *(Ornithorhynchus anatinus*): a fine-scale look at adjacent river systems. Australian Journal of Zoology, 57(4):225–234.

Kong, A., Frigge, M. L., Masson, G., Besenbacher, S., Sulem, P., Magnusson, G., Gudjonsson, S. A., Sigurdsson, A., Jonasdottir, A., Jonasdottir, A., Wong, W. S. W., Sigurdsson, G., Walters, G. B., Steinberg, S., Helgason, H., Thorleifsson, G., Gudbjartsson, D. F., Helgason, A., Magnusson, O. T., Thorsteinsdottir, U., and Stefansson, K. (2012). Rate of de novo mutations and the importance of father’s age to disease risk. Nature, 488(7412):471–475.

Lamichhaney, S., Berglund, J., Almén, M. S., Maqbool, K., Grabherr, M., Martinez-Barrio, A., Promerova, M., Rubin, C.-J., Wang, C., Zamani, N., et al. (2015). Evolution of Darwin/’s finches and their beaks revealed by genome sequencing. Nature, 518(7539):371–375.

Lawson, D. J., Hellenthal, G., Myers, S., and Falush, D. (2012). Inference of population structure using dense haplotype data. PLoS Genet, 8(1):e1002453.

Leslie, S., Winney, B., Hellenthal, G., Davison, D., Boumertit, A., Day, T., Hutnik, K., Royrvik, E. C., Cunliffe, B., W. T. C. C. C., I. M. S. G. C., Lawson, D. J., Falush, D., Freeman, C., Pirinen, M., Myers, S., Robinson, M., Donnelly, P., and Bodmer, W. (2015). The fine-scale genetic structure of the British population. Nature, 519(7543):309–314.

Li, H. and Durbin, R. (2011). Inference of human population history from individual whole-genome sequences. Nature, 475(7357):493–496.

Li, M. X., Kwan, J. S., Bao, S. Y., Yang, W., Ho, S. L., Song, Y. Q., and Sham, P. C. (2013). Predicting mendelian disease-causing non-synonymous single nucleotide variants in exome sequencing studies. PLoS Genet, 9(1):e1003143.

Locke, D. P., Hillier, L. W., Warren, W. C., Worley, K. C., Nazareth, L. V., Muzny, D. M., Yang, S.-P., Wang, Z., Chinwalla, A. T., Minx, P., et al. (2011). Comparative and demographic analysis of orang-utan genomes. Nature, 469(7331):529–533.

Lunter, G. and Goodson, M. (2011). Stampy: a statistical algorithm for sensitive and fast mapping of Illumina sequence reads. Genome Res, 21(6):936–9.

Lunter, Gerton Goodson, Martin 075491/Z/04/Wellcome Trust/United Kingdom Genome Res. 2011 Jun;21(6):936–9. Epub 2010 Oct 27.

Malinsky, M., Challis, R. J., Tyers, A. M., Schiffels, S., Terai, Y., Ngatunga, B. P., Miska, E. A., Durbin, R., Genner, M. J., and Turner, G. F. (2015). Genomic islands of speciation separate cichlid ecomorphs in an East African crater lake. Science, 350(6267):1493–1498.

Manichaikul, A., Mychaleckyj, J. C., Rich, S. S., Daly, K., Sale, M., and Chen, W.-M. (2010). Robust relationship inference in genome-wide association studies. Bioinformatics, 26(22):2867–2873.

McManus, K. F., Kelley, J. L., Song, S., Veeramah, K. R., Woerner, A. E., Stevison, L. S., Ryder, O. A., Kidd, J. M., Wall, J. D., Bustamante, C. D., et al. (2015). Inference of gorilla demographic and selective history from whole-genome sequence data. Molecular biology and evolution, 32(3):600–612.

Miller, W., Schuster, S. C., Welch, A. J., Ratan, A., Bedoya-Reina, O. C., Zhao, F., Kim, H. L., Burhans, R. C., Drautz, D. I., Wittekindt, N. E., et al. (2012). Polar and brown bear genomes reveal ancient admixture and demographic footprints of past climate change. Proceedings of the National Academy of Sciences, 109(36):E2382–E2390.

Moss, P., Tibby, J., Petherick, L., McGowan, H., and Barr, C. (2012). Late Quaternary vegetation history of North Stradbroke Island, Queensland, eastern Australia. Quaternary Science Reviews, 74:257–272.

Musser, A. (1998). Evolution, biogeography and palaeoecology of the ornithorhynchidae. Australian Mammology, 20(2):147–162.

Musser, A. (2013). Review of the monotreme fossil record and comparison of palaeontological and molecular data. Comp. Biochem. Physiol., Part A Mol. Integr. Physiol., 136:927–942.

Nadachowska-Brzyska, K., Burri, R., Olason, P. I., Kawakami, T., Smeds, L., and Ellegren, H. (2013). Demographic divergence history of pied flycatcher and collared flycatcher inferred from whole-genome re-sequencing data. PLoS Genet, 9(11):e1003942.

Nadachowska-Brzyska, K., Burri, R., Smeds, L., and Ellegren, H. (2016). PSMC-analysis of effective population sizes in molecular ecology and its application to black-and-white Ficedula flycatchers. Molecular Ecology.

Nei, M. and Li, W.-H. (1979). Mathematical model for studying genetic variation in terms of restriction endonucleases. Proceedings of the National Academy of Sciences, 76(10):5269–5273.

Novembre, J. and Ramachandran, S. (2011). Perspectives on human population structure at the cusp of the sequencing era. Annu Rev Genomics Hum Genet, 12:245–274.

O’Connell, J., Gurdasani, D., Delaneau, O., Pirastu, N., Ulivi, S., Cocca, M., Traglia, M., Huang, J., Huffman, J. E., Rudan, I., McQuillan, R., Fraser, R. M., Campbell, H., Polasek, O., Asiki, G., Ekoru, K., Hayward, C., Wright, A. F., Vitart, V., Navarro, P., Zagury, J.-F., Wilson, J. F., Toniolo, D., Gasparini, P., Soranzo, N., Sandhu, M. S., and Marchini, J. (2014). A General Approach for Haplotype Phasing across the Full Spectrum of Relatedness. PLoS Genetics, 10(4):e1004234.

Patterson, N., Price, A. L., and Reich, D. (2006). Population structure and eigenanalysis. PLoS Genet, 2(12):e190.

Phillips, S. J., Anderson, R. P., and Schapire, R. E. (2006). Maximum entropy modeling of species geographic distributions. Ecological modelling, 190(3):231–259.

Pian, R., Archer, M., and S.J., H. (2013). A new, giant platypus, obdurodon tharalkooschild, sp. nov. (monotremata, ornithorhynchidae), from the riversleigh world heritage area, australia. Journal of Vertebrate Paleontology, 33(6):1255–1259.

Prado-Martinez, J., Sudmant, P. H., Kidd, J. M., Li, H., Kelley, J. L., Lorente-Galdos, B., Veeramah, K. R., Woerner, A. E., O’Connor, T. D., Santpere, G., Cagan, A., Theunert, C., Casals, F., Laayouni, H., Munch, K., Hobolth, A., Halager, A. E., Malig, M., Hernandez-Rodriguez, J., Hernando-Herraez, I., Prüfer, K., Pybus, M., Johnstone, L., Lachmann, M., Alkan, C., Twigg, D., Petit, N., Baker, C., Hormozdiari, F., Fernandez-Callejo, M., Dabad, M., Wilson, M. L., Stevison, L., Camprubí, C., Carvalho, T., Ruiz-Herrera, A., Vives, L., Mele, M., Abello, T., Kondova, I., Bontrop, R. E., Pusey, A., Lankester, F., Kiyang, J. A., Bergl, R. A., Lonsdorf, E., Myers, S., Ventura, M., Gagneux, P., Comas, D., Siegismund, H., Blanc, J., Agueda-Calpena, L., Gut, M., Fulton, L., Tishkoff, S. A., Mullikin, J. C., Wilson, R. K., Gut, I. G., Gonder, M. K., Ryder, O. A., Hahn, B. H., Navarro, A., Akey, J. M., Bertranpetit, J., Reich, D., Mailund, T., Schierup, M. H., Hvilsom, C., Andrés, A. M., Wall, J. D., Bustamante, C. D., Hammer, M. F., Eichler, E. E., and Marques-Bonet, T. (2013). Great ape genetic diversity and population history. Nature, 499(7459):471–475.

Pritchard, J. K., Stephens, M., and Donnelly, P. (2000). Inference of population structure using multilocus genotype data. Genetics, 155(2):945–959.

Purcell, S., Neale, B., Todd-Brown, K., Thomas, L., Ferreira, M. A. R., Bender, D., Maller, J., Sklar, P., de Bakker, P. I. W., Daly, M. J., and Sham, P. C. (2007). PLINK: a tool set for whole-genome association and population-based linkage analyses. Am J Hum Genet, 81(3):559–575.

Rimmer, A., Phan, H., Mathieson, I., Iqbal, Z., Twigg, S. R. F., Consortium, W., Wilkie, A. O. M., McVean, G., and Lunter, G. (2014). Integrating mapping-, assembly- and haplotype-based approaches for calling variants in clinical sequencing applications. Nat Genet, 46(8):912–918.

Rosauer, D., Catullo, R., VanDerWall, J., Moussalli, A., and Moritz, C. (2015). Lineage Range Estimation Method Reveals Fine-Scale Endemism Linked to Pleistocene Stability in Australian Rainforest Herpetofauna. PLoS One.

Schäuble, C. and Moritz, C. (2001). Comparative phylogeography of two open forest frogs from eastern Australia. Biological Journal of the Linnean Society, 74(2):157–170.

Serena, M. & Williams, G. (2012). Movements and cumulative range size of the platypus (Ornithorhynchus anatinus) inferred from mark?recapture studies. Australian Journal of Zoology.

Serena, M., Williams, G., Weeks, A., and J., G. (2014). Variation in platypus (Ornithorhynchus anatinus) life-history attributes and population trajectories in urban streams. Australian Journal of Zoology, 62(3):223–234.

Thomas, C. G., Wang, W., Jovelin, R., Ghosh, R., Lomasko, T., Trinh, Q., Kruglyak, L., Stein, L. D., and Cutter, A. D. (2015). Full-genome evolutionary histories of selfing, splitting, and selection in Caenorhabditis. Genome Res, 25(5):667–678.

Uchimura, A., Higuchi, M., Minakuchi, Y., Ohno, M., Toyoda, A., Fujiyama, A., Miura, I., Wakana, S., Nishino, J., and Yagi, T. (2015). Germline mutation rates and the long-term phenotypic effects of mutation accumulation in wild-type laboratory mice and mutator mice. Genome Research, 25(8):1125–1134.

Venn, O., Turner, I., Mathieson, I., de Groot, N., Bontrop, R., and McVean, G. (2014). Strong male bias drives germline mutation in chimpanzees. Science, 344(6189).

Wallberg, A., Han, F., Wellhagen, G., Dahle, B., Kawata, M., Haddad, N., Simões, Z. L. P., Allsopp, M. H., Kandemir, I., De la Rúa, P., Pirk, C. W., and Webster, M. T. (2014). A worldwide survey of genome sequence variation provides insight into the evolutionary history of the honeybee Apis mellifera. Nat Genet, 46(10):1081–1088.

Warren, W. C., Hillier, L. W., Marshall Graves, J. A., Birney, E., Ponting, C. P., Grutzner, F., Belov, K., Miller, W., Clarke, L., Chinwalla, A. T., Yang, S. P., Heger, A., Locke, D. P., Miethke, P., Waters, P. D., Veyrunes, F., Fulton, L., Fulton, B., Graves, T., Wallis, J., Puente, X. S., Lopez-Otin, C., Ordonez, G. R., Eichler, E. E., Chen, L., Cheng, Z., Deakin, J. E., Alsop, A., Thompson, K., Kirby, P., Papenfuss, A. T., Wakefield, M. J., Olender, T., Lancet, D., Huttley, G. A., Smit, A. F., Pask, A., Temple-Smith, P., Batzer, M. A., Walker, J. A., Konkel, M. K., Harris, R. S., Whittington, C. M., Wong, E. S., Gemmell, N. J., Buschiazzo, E., Vargas Jentzsch, I. M., Merkel, A., Schmitz, J., Zemann, A., Churakov, G., Kriegs, J. O., Brosius, J., Murchison, E. P., Sachidanandam, R., Smith, C., Hannon, G. J., Tsend-Ayush, E., McMillan, D., Attenborough, R., Rens, W., Ferguson-Smith, M., Lefevre, C. M., Sharp, J. A., Nicholas, K. R., Ray, D. A., Kube, M., Reinhardt, R., Pringle, T. H., Taylor, J., Jones, R. C., Nixon, B., Dacheux, J. L., Niwa, H., Sekita, Y., Huang, X., Stark, A., Kheradpour, P., Kellis, M., Flicek, P., Chen, Y., Webber, C., Hardison, R., Nelson, J., Hallsworth-Pepin, K., Delehaunty, K., Markovic, C., Minx, P., Feng, Y., Kremitzki, C., Mitreva, M., Glasscock, J., Wylie, T., Wohldmann, P., Thiru, P., Nhan, M. N., Pohl, C. S., Smith, S. M., Hou, S., Nefedov, M., et al. (2008). Genome analysis of the platypus reveals unique signatures of evolution. Nature, 453(7192):175–83.

Weber, L., VanDerWall, J., Schmidt, S., McDonald, W., and L.P., S. (2014). Patterns of rain forest plant endemism in subtropical Australia relate to stable mesic refugia and species dispersal limitations. Journal of Biogeography, 41:222–238.

Welch, J. J., Bininda-Emonds, O. R., and Bromham, L. (2008). Correlates of substitution rate variation in mammalian protein-coding sequences. BMC Evolutionary Biology, 8(1):53.

Xue, Y., Prado-Martinez, J., Sudmant, P. H., Narasimhan, V., Ayub, Q., Szpak, M., Frandsen, P., Chen, Y., Yngvadottir, B., Cooper, D. N., de Manuel, M., Hernandez-Rodriguez, J., Lobon, I., Siegismund, H. R., Pagani, L., Quail, M. A., Hvilsom, C., Mudakikwa, A., Eichler, E. E., Cranfield, M. R., Marques-Bonet, T., Tyler-Smith, C., and Scally, A. (2015). Mountain gorilla genomes reveal the impact of long-term population decline and inbreeding. Science, 348(6231):242–245.

Zhan, S., Zhang, W., Niitepõld, K., Hsu, J., Haeger, J. F., Zalucki, M. P., Altizer, S., De Roode, J. C., Reppert, S. M., and Kronforst, M. R. (2014). The genetics of monarch butterfly migration and warning colouration. Nature, 514(7522):317–321.

